# Single step, high efficiency CRISPR-Cas9 genome editing in primary human disease-derived fibroblasts

**DOI:** 10.1101/440099

**Authors:** Matteo Martufi, Robert B. Good, Radu Rapiteanu, Tobias Schmidt, Eleni Patili, Ketil Tvermosegaard, Carmel B. Nanthakumar, Joanna Betts, Andy D. Blanchard, Klio Maratou

## Abstract

Genome editing is a tool that has many applications including the validation of potential drug targets. However, performing genome editing in low passage, primary human cells with the greatest physiological relevance, is notoriously difficult. High editing efficiency is desired because it enables gene knock outs (KO) to be generated in bulk cellular populations and circumvents the problem of having to generate clonal cell isolates. Here, we describe a single step workflow enabling >90% KO generation in primary human lung fibroblasts via CRISPR ribonucleoprotein delivery, in the absence of antibiotic selection or clonal expansion. As proof of concept, we performed a disease relevant phenotypic assay measuring collagen deposition in response to TGFβ and demonstrated SMAD3 but not SMAD2 dependent deposition of type I collagen following knockout of each using our single step methodology. The optimization of this workflow can readily be transferred to other primary cell types.

## Introduction

One of the remaining challenges for genome editing is to perform experiments in primary cells, isolated from patient or healthy donor tissues and used experimentally at low passages to minimise cell changes in culture. The most widely used workflows for genome editing involve monoclonal cell isolation prior to subsequent characterisation of the effect of the edit. The generation of clonal cells ensures that phenotypic experiments are performed using a uniform, genetically identical population of cells. However, primary cells cannot proliferate indefinitely or survive outside of specific culture conditions, and therefore are not amenable to monoclonal selection, or clonal expansion following genome editing. One solution is to use the pool of edited cells (bulk cell culture) directly for experimental analysis. In this case, the editing efficiency needs to be sufficiently high, so that so that a large proportion if not all cells contain the desired modification at all copies of the target locus. Such analysis is suited for functional analysis of genes and pathways, as it accelerates the timelines for validation of novel targets and leads to a better understanding of the biological mechanisms underlying human diseases.

To develop genome editing workflows in human primary cells we chose to focus on primary human lung fibroblasts which are important for the study of molecular pathways involved in Idiopathic Pulmonary Fibrosis (IPF). Patients with IPF have a poor prognosis, with median survival of 3 years post diagnosis, and a progressive loss of lung function due to the synthesis and deposition of a local, dense collagen-rich extracellular matrix (ECM)^1^. Understanding the mechanisms under-pinning ECM secretion and deposition has important therapeutic implications and therapeutic approaches targeting these mechanisms are being explored clinically. The ability to rapidly and effectively knockout individual genes in freshly isolated cells from patients would provide a valuable early target validation platform to assess novel mechanistic approaches.

Accurate genotyping of the edited cells is an important requirement for bulk cell culture editing pipelines. It confirms on-target editing and provides precise measurements of the editing events. Most commonly, genotyping is achieved by Surveyor nuclease^2^, T7 endonuclease I (T7E1) assay^3^, TIDE assay^4^, or droplet digital PCR^5^. These methods are low throughput, cannot be easily multiplexed and do not provide accurate sequence information on the achieved edits. Moreover, they can’t be easily used to genotype a bulk population of cells with several different mutations. The development of workflows that use targeted deep sequencing ^6,7,8,9,10^ has solved this problem and paved the way for automated, target focused genome editing at scale. Our lab adopted the publicly available sequence-evaluation tool OutKnocker^6,7^, which allows rapid identification of all-allelic frameshift mutations in bulk cellular populations.

Here we describe how we established a CRISPR/Cas9 ribonucleoprotein (RNP) complex workflow to carry out highly efficient genome editing in a bulk population of primary fibroblasts derived from IPF patients without applying any selection. To optimise the electroporation of RNP complex delivery into fibroblasts, we edited gene PI4KA and established conditions enabling full gene KO in bulk cells with a single round of electroporation. Using these conditions, we could replicate results with multiple targets and we present SMAD2 and SMAD3 single KOs, as well as a double KO, as a proof of concept. The pipeline described in this paper is presented as a tool that can be applied in target validation studies for drug discovery in allowing the rapid and efficient genomic modification of any gene and further opens the possibility to identify associated clinical biomarkers.

## Results

### Optimisation of workflow for genome editing in primary human lung fibroblasts

We hypothesise that phenotypic assays could be performed in bulk populations with over 90% of alleles for a gene of interest containing indel mutations since this would translate into a very low chance of obtaining a cell carrying two wild type alleles. To achieve this, we base our experiments on RNP delivery using electroporation. We built our workflow using the commercially available Alt-R CRISPR-Cas9 system (IDT) which comprises of a chemically-modified crisprRNA (crRNA) and trans-activating crisprRNA (tracrRNA) complexed with Cas9 protein.

To fine-tune our workflow, we focused on generating KO cells for Phosphatidylinositol 4-kinase alpha (PI4KA), a moderately expressed cell signalling protein. A critical factor in successful gene editing is the choice of highly efficient crRNAs. We first carried out screening of eight crRNAs targeting different exons of PI4KA, to identify the two best performing ones (figure S1). The RNP complex was delivered by electroporation using a Nucleofector 2b device with manufacturer (Lonza) recommended electroporation conditions (figure S1). Primary fibroblasts were left in culture for 24 hours, before being collected and genotyped through targeted deep sequencing using an Illumina MiSeq system^7^. Analysis of the generated mutations was carried out with the freely available web tool OutKnocker^6^, which provides detailed information on the type and incidence of mutations in the allele population (Figure 1; stage 1). Despite screening for multiple crRNAs, we observed that editing efficiency was too low to enable functional analysis of the bulk cellular population (figure S1). Therefore, we complemented the crRNA selection with a thorough optimization of the electroporation delivery step.

**Figure 1.**
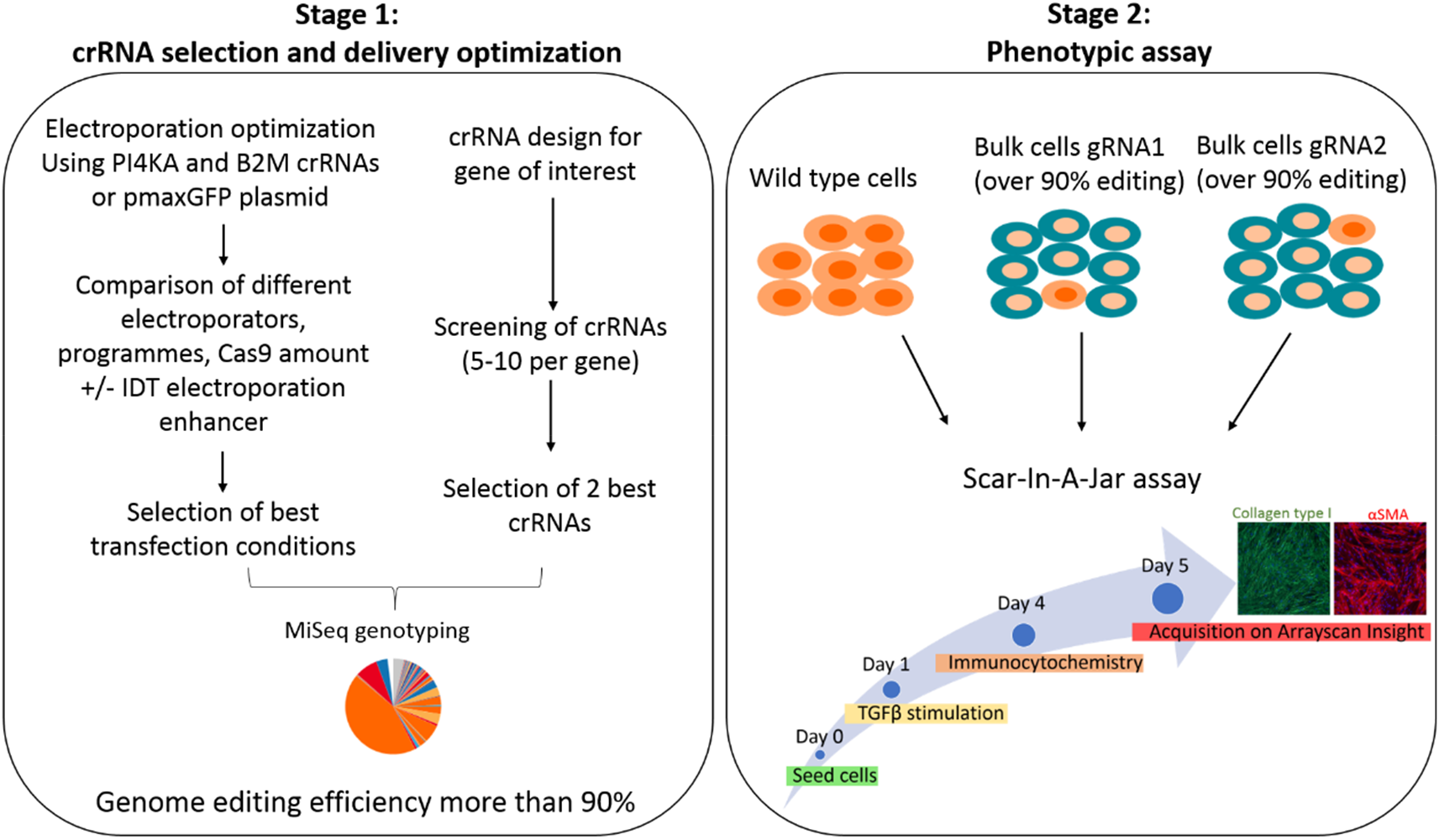
Pipeline of genome editing in primary human lung fibroblasts. Stage1: Parameters such as different Nucleofector systems, electroporation programs, Cas9 amounts, electroporation buffers and an electroporation enhancer are tested to find the best condition in terms of cell morphology, cell viability and genome editing efficiency. gRNA screening of 5-10 per gene is carried out to select the two best gRNAs. Genotyping of the bulk cell population to quantify the editing efficiency is performed by MiSeq sequencing and OutKnocker analysis. Pie chart represents alleles in the cell population, with distinct colours for wild type sequences, in-frame indels, or out-of-frame indels. Stage 2: Bulk cell populations with more than 90% indels are used to perform the phenotypic assay Scar-in-aJar. This assay measures type I collagen deposition and the αSMA expression upon TGFβ stimulation.

We made a side by side comparison of the Lonza Nucleofector 2b and 4D systems. For the 2b system we utilised a Basic Nucleofector kit for Primary Fibroblasts and the already established and manufacturer recommended program A-24. However, this could not be transferred to the 4D system and therefore we proceeded with testing several electroporation programs via transfecting cells with the positive control plasmid pmaxGFP, which is included in each Nucleofector™ Kit (figure S2A), and the P3 Primary Cell 4D-Nucleofector™ X Kit. Three electroporation programs (CA137, CM137 and CM138) were selected on the basis of transfection efficiency and cell morphology. To find the best program, an RNP complex, formed using 15pmol Cas9, with 1.2:1 RNA/Cas9 ratio, was delivered to the cells targeting exon 1 of the B2M gene, which has previously been successfully used in other cell types^11,12^ (figure S2B). Alongside optimization of the electroporation program, we tested the benefits of co-transfection with the Alt-R Cas9 electroporation enhancer (IDT). Inclusion of the enhancer dramatically increased indel generation (figure S2B). Genome editing efficiency was similar among the three programs, however we decided to use the Nucleofector 4D program CM138 due to better cell recovery after electroporation. With the B2M guide we only achieve approximately 45% editing efficiency. However, for this target we used a guide optimised for other cell types, where high efficiency was not required. To proceed with final optimisations, we returned to use of the PI4KA exon 10 guide RNA selected after the initial guide RNA screening described above (figure S1). This time we increased the amount of Cas9 to 60pmol per reaction. For Nucleofector 2b we could not achieve very high editing efficiency only with one electroporation, and therefore had to perform another round of electroporation to get to indel percentages that would allow us to perform phenotypic assays in a bulk population. Contrarily, for Nucleofector 4D, a single round of electroporation in presence of the electroporation enhancer using program CM138 with 60pmol Cas9 in a ratio of 1.2:1 RNA/Cas9, was enough to achieve 100% editing efficiency (figure 2B). This condition was used to perform all remaining experiments described in this paper.

**Figure 2.**
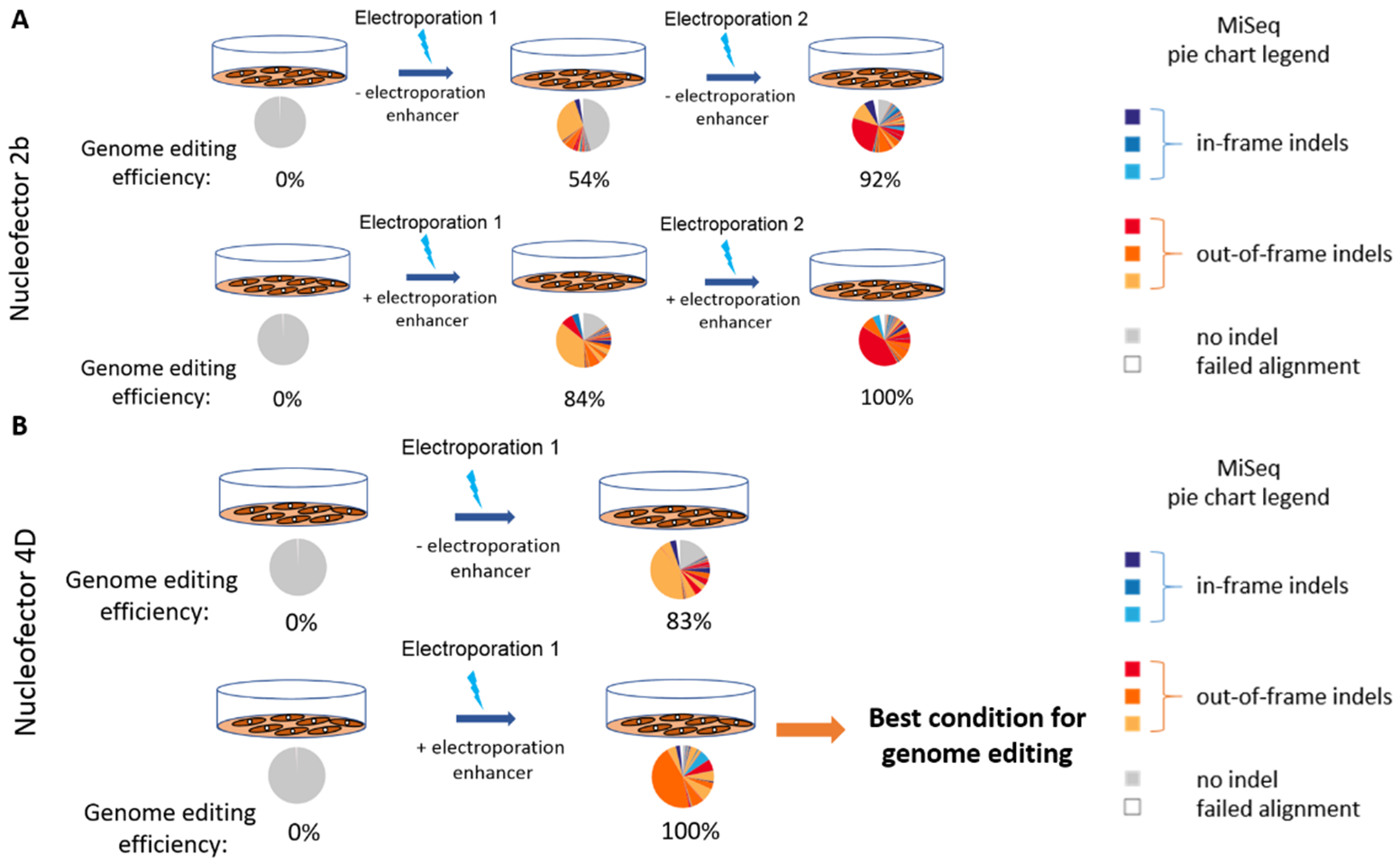
PI4KA genome editing optimization. Comparison between the Nucleofector 2b and the 4D was carried out by using a gRNA targeting exon 10 of PI4KA and experiments were performed in presence and absence of an electroporation enhancer. Cells were collected after 24 hours after electroporation and genotyped by MiSeq and OutKnocker web tool analysis. A pie chart represents the OutKnocker analysis output. Every pie chart represents a cellular pool, whilst colours of the individual pie areas indicate in-frame mutations (blue), out-of-frame mutations (red), or no indel calls (grey). **A)** For Nucleofector 2b, two rounds of electroporation are required to achieve high genome editing efficiency, even in presence of an electroporation enhancer. **B)** For Nucleofector 4D, one round of electroporation is sufficient for high editing efficiency.

### SMAD3 KO affects the fibrotic response to TGFβ stimulation

Once cells have been edited with high efficiency, they can be assayed within a disease relevant phenotypic assay. We tested for collagen type I deposition and alpha smooth muscle actin (αSMA) expression driven by TGFβ stimulation and macromolecular crowding, using a high content imaging assay termed Scar-in-a-Jar (Stage 2, figure 1)^13,14^. This phenotypic, high content screening method quantifies the amount of extracellular collagen deposition to deliver a robust and reliable end-point to study the effect of gene ablation in fibrosis.

To demonstrate the utility of our workflow in a setting relevant to fibrosis, we knocked-out two closely related proteins critical to TGFβ signalling (SMAD2 and SMAD3). These are two transcription factors involved in the activation of the fibrotic response and are responsible for the upregulation of genes such as collagen type I and αSMA^15^. Seven gRNAs targeting different exons were tested for each gene (sequence shown in supplementary Table S1). The exon 6 SMAD3 gRNA, showing nearly 100% mutation efficiency, alongside the exon 6 SMAD2 gRNA, exhibiting nearly 80% mutation efficiency, were selected for progression to phenotypic experiments using the Scar-in-a-Jar assay (figure 3A). The exon 6 SMAD2 gRNA performed with around 94% editing efficiency when the experiment was subsequently repeated (figure 3B; Donor 1 n=4, Donor 2 n=3). The SMAD3 exon 2, exon 4 and exon 6 guides also showed consistently high editing efficiencies with experimental repeats (figure 3B), providing us with high confidence in our workflow. The high efficiency in generating mutations in the two gene loci is mirrored by absence of the proteins. Although SMAD2 and SMAD3 proteins share more than 90% of protein sequence homology, western blot analysis confirmed that each KO was protein specific (figure 4A).

**Figure 3.**
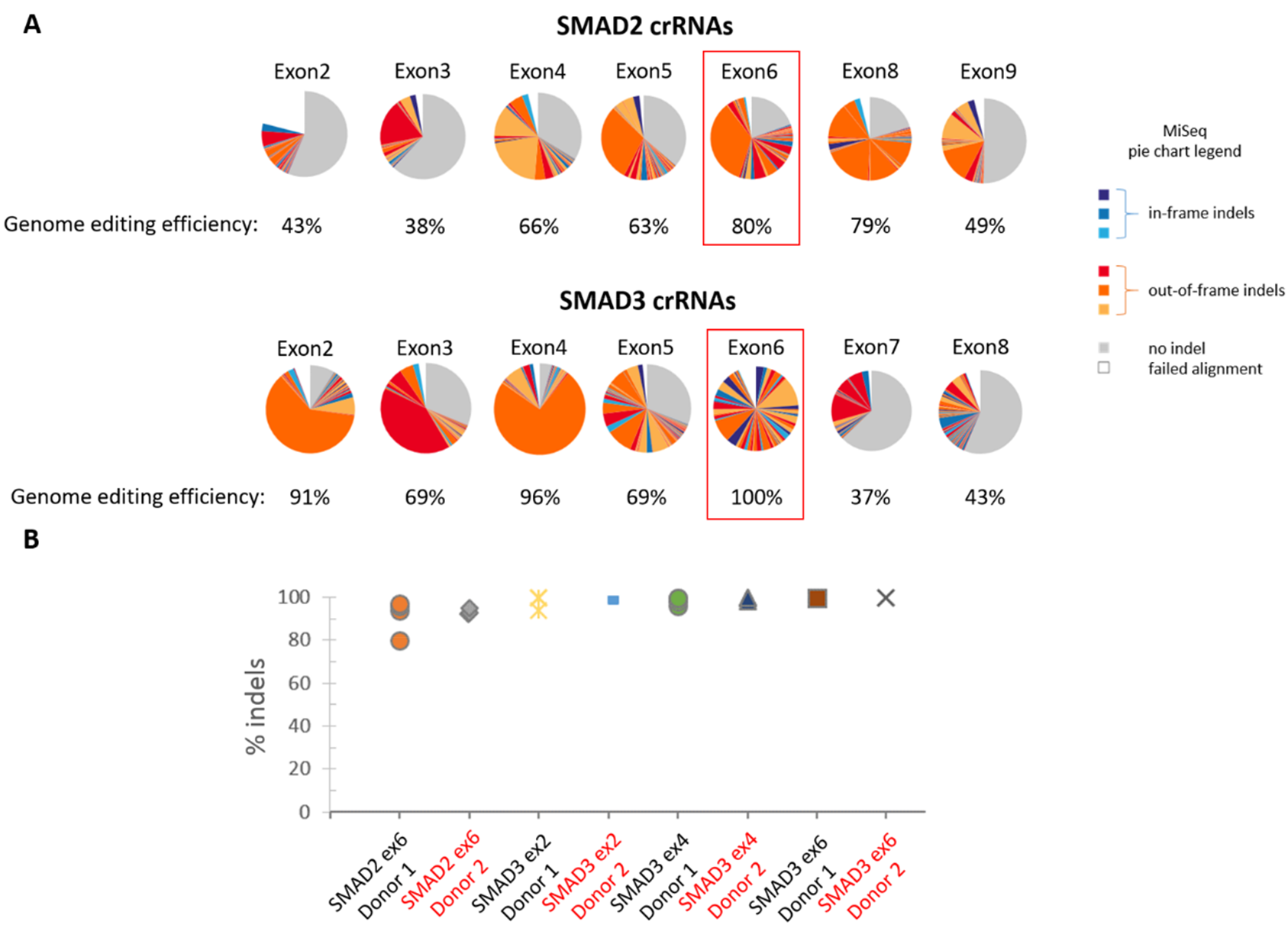
Efficient genome editing for SMAD2 and SMAD3. **A)** Pie charts represent the OutKnocker genotyping analysis results from one representative experiment, and illustrate the mutations generated by gRNAs targeting different exons for SMAD2 and SMAD3. The crRNAs highlighted were chosen to edit cells in subsequent experiments. **B)** Percentage of indels measured in all experimental and technical replicates, using one gRNA for SMAD2 and three for SMAD3, in two donors (Donor 1 n=4, Donor 2 n=3).

**Figure 4.**
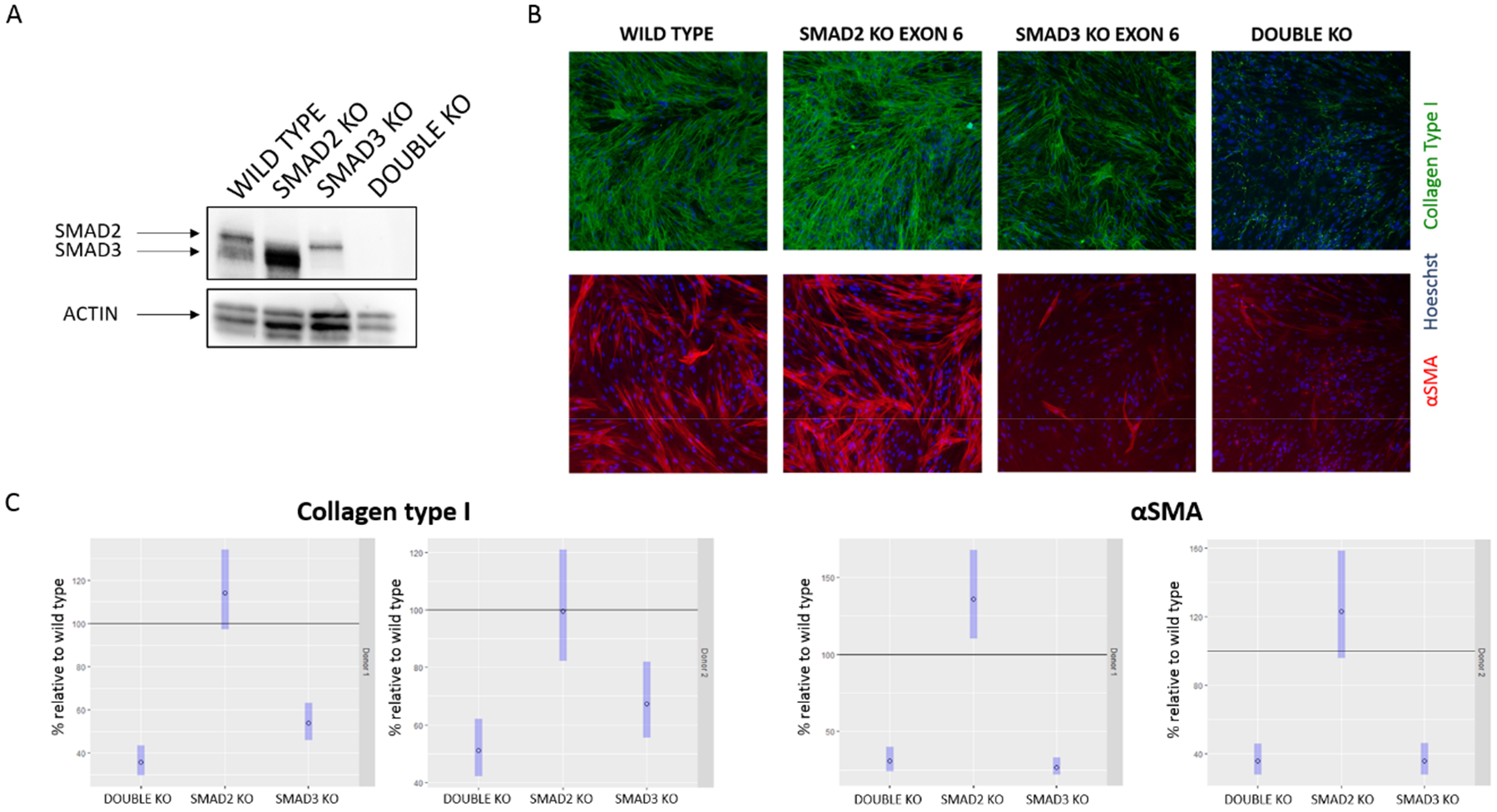
Scar-in-a-Jar shows collagen deposition reduction in SMAD3 KO and double KO cells. **A)** Western blot analysis from wild type, SMAD2 KO, SMAD3 KO and double SMAD2/SMAD3 KO cells. One antibody recognising both SMAD proteins was used as the two proteins share more than 90% homology of amino acid sequence. ACTIN was the loading control. Full length western blot is presented in Supplementary figure 5. **B)** Scar-in-a-Jar assay results. Collagen deposition was measured by immunofluorescence, cells were also stained for Hoechst to visualize the nuclei and αSMA to detect the level of fibroblast differentiation. **C)** Scar-in-a-Jar quantification of fluorescence estimated contrasts between KOs and wildtype TGFβ-stimulated cells, across donors. The plots show the collagen type I and αSMA results (Donor 1 n=3, Donor 2 n=2). Dots are estimated differences; blue bars are 95% confidence intervals.

In a separate experiment, a double knockout of both proteins in the same cell population was performed. We chose the exon6 gRNAs for both SMAD2 and SMAD3 and co-delivered the RNP complexes during the same electroporation. The experiment was performed in two donors. Genotyping of both loci showed high efficient genome editing for both genes in the same sample (figure S4). This was confirmed by western blot that showed ablation of both proteins (figure 4A).

Using the exon 6 gRNAs targeting SMAD3 andSMAD2, we performed Scar-in-a-Jar assays, in two different donors (n = 3 for donor 1; n = 2 for donor 2). KO of SMAD3 resulted in a dramatic reduction in collagen deposition in comparison to the wild type IPF fibroblasts, while KO of SMAD2 did not show any reduction (figure 4B, 4C and S3). We checked the levels of αSMA as an additional phenotypic measurement. This gene is upregulated when fibroblasts undergo myofibroblast differentiation after TGFβ stimulation and is a crucial requirement to enable cells to produce collagen^16^. The SMAD3 KO cells lacked αSMA expression, while the SMAD2 KOs showed increased expression in comparison to wild type cells (figure 4B, 4C and S3), supporting the result obtained for collagen deposition. Furthermore, in the double KO, we observed a decrease in collagen deposition as well as lack of expression of αSMA (figure 4B, 4C and S3). These results differentiate the two SMAD proteins, demonstrating a critical role of SMAD3 in response to TGFβ stimulation for collagen production, in contrast to SMAD2, in the Scar-in-a-Jar assay.

## Discussion

Selection of disease relevant targets is an important initial step in the drug discovery process. CRISPR technology enables perturbation of genes and pathways in human disease relevant cellular systems as well as model organisms, enabling better understanding of the function and biology of a target^17^. However, the majority of published work utilising gene editing is performed in less physiologically relevant immortalised cell lines, because genome editing in primary cells presents numerous technical challenges. Here, we sought to optimise a gene editing workflow in patient-derived primary lung fibroblasts that enables studies of key mechanisms involved in lung fibrosis. We developed and extensively optimised a pipeline that uses easily accessible CRISPR/Cas9 RNP reagents and Nucleofector 4D electroporation, which results in over 90% KO without the use of antibiotic selection or monoclonal expansion, reducing culture time. Moreover, our protocol allows the production of double gene KO cells as demonstrated by the ablation of SMAD2 and SMAD3 proteins within the same sample. This result shows an important advancement of genome editing technology in primary cells and we deliver a method that can be used to study the combinatorial effect of genes in a disease context.

Other groups have also reported genome editing in primary human cells, such as: lung or dermal fibroblasts^18^, airway epithelia^18,19^, endothelial cells^20^, albeit at a reduced editing efficiency. These studies rely on the use of plasmid or viral CRISPR vectors. CRISPR delivered as RNP complex has multiple advantages over these other formats, such as rapid editing within 3 hours and fast clearance from cells by 24 hours^21^, minimising the possibility of off-target effects. Recently, highly efficient genome editing was achieved in primary human and mouse T cells^22,23^, as well as CD34+ hematopoietic stem and progenitor cells ^23^, by using RNP electroporation and a protocol similar to ours. Our workflow adds the important contribution of deep sequencing, which enables accurate editing efficiency measurement regardless of cellular localization of the targeted proteins. Moreover, in our workflow electroporation is performed using sample strips that process 16 different gRNAs simultaneously. Coupled with the deep sequencing genotyping, which measures hundreds of samples at the same time, we are able to increase the throughput of editing, enabling the generation of multiple gene KOs in one experiment. This can be critical in a context of target validation for a specific pathway or for a list of genes connected to a disease, because it allows to study the function of many genes in the same experiment.

Optimisation of bespoke editing conditions is required for every primary cell type used in an experiment. The process to identify optimum conditions used in this paper is relevant to any type of cell line or tissue. Achievement of near 100% editing efficiency will require identification of guides that cut with high efficiency and juxtaposition with optimum transfection conditions. To transfect primary fibroblasts, we found that electroporation with the 4D Nucleofector in the presence of an IDT electroporation enhancer gives the best results. We reproducibly achieved over 90% editing efficiencies with over 10 targets (data not shown). It is worth noting though, that some primary cell types, e.g. macrophages, do not tolerate inclusion of the electroporation enhancer. Other parameters that are critical and need investigation are different electroporation buffers, cell density optimisation and examination of different gRNA-to-Cas9 ratios.

Our workflow was developed to study targets for lung fibrosis. To demonstrate the value of our pipeline, we performed genome editing of SMAD2 and SMAD3 proteins which function as transcriptional modulators activated by transforming growth factor-beta. The signalling cascade activated by TGFβ triggers the phosphorylation of each protein that then form a complex with SMAD4. The SMAD proteins complexed together enter the nucleus and promote the transcription of pro-fibrotic genes^24^. However, it has been shown that they may be responsible for activation of different transcriptional programs^24–26^. In the context of disease, SMAD3 but not SMAD2 appears to be important for mediating TGFβ signalling in renal fibrosis^27^, hepatic fibrosis^28^ and cardiac fibrosis^29^. In contrast, SMAD2 is crucial for epiblast development and patterning of three germ layers during early developmental events^25^. In agreement with the published data for other types of fibrosis, we also show in the context of IPF reduced amount of deposited collagen and lack of the myofibroblast differentiation marker αSMA in SMAD3 KO, but not SMAD2 KO, upon TGFβ stimulation. Moreover, in SMAD2 KO we observed an increased expression of αSMA probably due to an enhanced activation of SMAD3 as already shown in hepatic and renal fibrosis^27,28^. This indicates that the presence of SMAD3 is critical for cell differentiation into myofibroblasts and collagen production in primary disease fibroblasts.

In this paper we have tested the ability of cells to differentiate and produce collagen by performing a high content cell imaging assay. Other phenotypic assays adopting transcriptomic analysis or measurement of secreted fibrosis mediators could also be deployed to examine what effect the perturbation of a target has in the context of lung fibroblast biology. It is worth noting that our workflow can be used to study the role of genes involved in several different pathologies in which fibroblasts from diverse organs contribute to tissue re-modelling and could be used as a primary target validation tool for novel fibrosis targets. Moreover, the optimisation process we have described to enable efficient genome editing can be applied to other primary cell types and hence should facilitate genome editing in other contexts. In summary, we describe a novel pipeline to study gene function in the context of lung fibrosis and deliver key considerations that will be useful for similar functional studies in other cell types.

## Methods

### Study approval

Samples of IPF lung tissue were obtained from patients undergoing lung transplant or surgical lung biopsy following informed signed consent and with research ethics committee approval (11/NE/0291, 10/H0504/9, 10/H0720/12 and 12/EM/0058). Lung tissues were obtained from Newcastle Upon Tyne Hospitals NHS Foundation Trust and no tissues were procured from prisoners. The human biological samples were sourced ethically, and their research use was in accord with the terms of the informed consents under an IRB/EC approved protocol. All experiments were performed in accordance with relevant guidelines and regulations.

### Cell culture

Primary human lung fibroblasts were grown from explant culture of IPF lung tissue as described previously^30^. Cells were cultured in Dulbecco’s Modified Eagle Medium (DMEM) (Gibco, #21969) supplemented with 10% heat inactivated fetal bovine serum and Glutamax. Cells were kept in a humidified 10% CO^2^ atmosphere at 37°C. Fibroblast vials were thawed at passage 2 and electroporated with CRISPR/Cas9 RNP complex two days after.

### CRISPR/Cas9 RNP electroporation

SMAD2, SMAD3 and PI4KA crRNA design was achieved using the free online tool Deskgen (https://www.deskgen.com/landing/). The crRNA sequences are reported in supplementary table 1. Alt-R crRNAs and Alt-R tracrRNA were acquired from IDT and re-suspended in nuclease-free duplex buffer (IDT) at a concentration of 100 μM. 1 μl from each of the two RNA components was mixed together and diluted in nuclease-free duplex buffer at a concentration of 25 μM. The mix was boiled at 95°C for 5 minutes and cooled down at room temperature for 10 minutes. 2.9 μl of the annealed RNA, corresponding to 72.5 pmoles, were complexed with 60 pmoles of Cas9 to a ratio of 1.2:1, unless stated otherwise. The mix was left at room temperature for 10 minutes. Afterwards, 60 pmol of electroporator enhancer (IDT) was added and incubated at room temperature with the RNP complex for 5 minutes.

Passage 2 cells from IPF donors were electroporated when around 80% confluent. For the 4D Nucleofector System (Lonza) experiments, 250,000 cells in 20 μl were used per electroporation. Cells were washed once in PBS, at 90g, 10 minutes, at room temperature. After PBS removal, cells were re-suspended in 15.5 μl of P3 solution from a P3 Primary Cell 4D-Nucleofector^®^ X Kit (Lonza) and mixed together with 4.5 μl of RNP complex to reach a total volume of 20 μl. Electroporation was performed in 16-well Nucleocuvette™ Strips, using program CM-138. For a double KO generation using the 4D Nucleofector system, RNPs targeting SMAD2 exon 6 and SMAD3 exon 6 were complexed in vitro as described above. 250,000 cells were washed once in PBS, at 90g, 10 minutes, at room temperature. After PBS removal, cells were re-suspended in 11 μl of P3 solution and mixed together with 9 μl of RNPs to reach a final volume of 20 μl. Electroporation was performed in 16-well Nucleocuvette™ Strips, using program CM-138. For the 2b Nucleofector System (Lonza) experiments, 1 million cells in 100 μl were used alongside a Basic Nucleofector™ kit for Primary Mammalian Fibroblasts (Lonza). After PBS removal, cells were re-suspended in 90 μl Basic Nucleofector™ Solution for Mammalian Fibroblasts.

### MiSeq genotyping analysis

Genotyping was carried out as described by Schmidt and colleagues^6^. All PCR primers are illustrated in supplementary tables 2 and 3. Briefly, 24 hours after electroporation, 10,000 cells were collected and resuspended in 10 μl of Lysis Buffer (0.2 mg/mL proteinase K, 1 mM CaCl2, 3 mM MgCl2, 1 mM EDTA, 1 % Triton X-100, 10 mM Tris (pH 7.5). Cells were incubated for 10 minutes at 65 °C and 15 minutes at 95 °C to generate a cell lysate and 2 μl of lysate were added in a first PCR reaction to amplify the genomic locus that flanks the CRISPR target site. A second PCR, using 1 μl of first PCR product, attaches Illumina adaptor and barcode sequence for sequencing and later deconvolution. For all PCRs, Phusion High-Fidelity Master Mix with HF Buffer (ThermoFisher) and 63°C annealing temperature were used. After the second PCR, samples were pooled and purified using AMPure XP beads (Beckman Coulter Life Sciences) in a ratio beads/PCR product 0.8:1. After purification, the sequencing library was quantified using a nanodrop instrument (ThermoFisher Scientific). 5 ng of library was denatured and diluted for NGS sequencing using a MiSeq instrument (Illumina) following the manufacturer′s instructions. Libraries were clustered and sequenced using 300bp single-end sequencing with a MiSeq Reagent Kit v2 (Illumina, MS-102-2002). Sequencing reads were analysed using the online tool OutKnocker v1.31 (http://www.OutKnocker.org/) using default parameters, as described by Schmid-Burgk and colleagues^6^.

### Scar-in-a-Jar

After genome editing, fibroblasts were left in culture for one week. Cells were then plated in a 96 well μ-clear imaging plates (BD Falcon) at a concentration of 1×10^4^ cells/well in DMEM (0.4% FBS). After 24 hours, cells were incubated in modified medium containing 0.4% of FBS, ascorbic acid (16.7μg/ml), Ficoll 70 (37.5mg/ml) and Ficoll 400 (25mg/ml) to generate macromolecular crowding conditions. TGFβ (1ng/ml) was also added and fibroblasts were left at 37°C for 72 hours before being fixed in ice-cold methanol and permeabilized in 0.1% Triton-X-100 in PBS. Immunostaining was performed by overnight incubation at 4°C with anti-collagen type I antibody (Sigma-Aldrich, C2456) at 1: 1,000 dilution in PBS. Cells were then incubated for one hour, at room temperature, with secondary antibody Alexa Fluor 488 goat anti-mouse IgG (Invitrogen, A11001) at 1:500 dilution and Hoeschst dye (Invitrogen H3570) at 1: 10,000 dilution in PBS. A further staining step with anti-αSMA antibody (Sigma C6198) at 1:1,000 dilution in PBS was carried out for 1 hour at room temperature. The culture plate was scanned using a CellInsight NXT HCS instrument (ThermoFisher) at 10x magnification.

### Scar-in-a-Jar results statistical analysis

Statistical analyses were performed in R. A linear mixed effects model was fitted to the data, using a fixed effect term corresponding to interaction between Genotype (SMAD2 KO, SMAD3 KO or Wildtype), TGFβ (plus or minus) and Donor (1 or 2). A random effect of Date was included, to account for day-to-day variations in the overall response level. The fit was performed using the R-package lme4^31^. The model described above was fitted separately to log2 of the Collagen response and log2 of the αSMA response. Because the interaction of genotype and TGFβ with Donor was statistically significant in both fits, the effect of knockout within TGFμ-stimulated cells was examined on a donor-by-donor basis, using Estimated Marginal Means obtained via the R-package emmeans^44^. To ease data visualisation in figure 4E, the estimated contrasts (KO-wildtype, within TGFμ-stimulated cells, for KO = SMAD2 KO and SMAD3 KO) and their confidence intervals were back-transformed from the log2 scale.

### Western blot

Protein extracts were produced using a cell pellet washed twice with PBS. 500,000 cells were lysed in 50 μl of RIPA buffer and left at 4°C for 30 minutes. The lysate was spun down at 14,000 rpm for 15 minutes and 20 μl of supernatant recovered and run on a 4-20% SDS-PAGE gel. Proteins were transferred onto a PVDF membrane using the iBLOT system (Invitrogen) for 6 minutes using program P3. The membrane was incubated with anti-SMAD 2/3 antibody (Cell Signalling Technology, #8685) at 1:1,000 dilution, or anti-Actin antibody (Sigma-Aldrich, A2228) as a loading control at 1:2,000 dilution, at 4°C overnight. Secondary goat anti-rabbit IgG HRP antibody (Invitrogen A16096) was used at 1:2,000 dilution for the anti-SMAD2/3 detections, whilst secondary rabbit anti-mouse IgG HRP antibody (Invitrogen A16160) was used at 1:5,000 dilution with anti-actin. Both secondary antibodies were incubated for 1 hour at room temperature. Visualization of membranes was performed using the machine Syngene G:BOX with the software Genesnap.

### Data availability

All data generated or analysed during this study are included in this published article (and its Supplementary Information files).

## Competing interests statement

The authors declare no competing interests.

## Acknowledgements

We would like to thank Bill Cairns and Quinn Lu for helpful comments and critical reading of the manuscript.

## Authors contribution

M.M led the experimental part, generated and analysed data; M.M and K.M drafted and edited the manuscript; M.M and R.B.G performed the Scar-in-a-Jar experiments; M.M and E.P performed the western blot experiments; K.T helped with experimental design and analysis of the Scar-in-a-Jar; R.R helped with electroporation optimization; T.S set up MiSeq genotyping; C.N set up Scar-in-a-Jar, J.B, A.D.B, K.M helped to conceive and design the study; K.M supervised the study. All the authors read and approved the final manuscript.

